# Optimal versus approximate channel selection methods for EEG decoding with application to topology-constrained neuro-sensor networks

**DOI:** 10.1101/2020.10.02.323501

**Authors:** Abhijith Mundanad Narayanan, Panagiotis Patrinos, Alexander Bertrand

## Abstract

Channel selection or electrode placement for neural decoding is a commonly encountered problem in electroencephalography (EEG). Since evaluating all possible channel combinations is usually infeasible, one usually has to settle for heuristic methods or convex approximations without optimality guarantees. To date, it remains unclear how large the gap is between the selection made by these approximate methods and the truly optimal selection. The goal of this paper is to quantify this optimality gap for several state-of-the-art channel selection methods in the context of least-squares based neural decoding. To this end, we reformulate the channel selection problem as a mixed-integer quadratic program (MIQP), which allows the use of efficient MIQP solvers to find the optimal channel combination in a feasible computation time for up to 100 candidate channels. As this reveals the exact solution to the combinatorial problem, it allows to quantify the performance losses when using state-of-the-art sub-optimal (yet faster) channel selection methods. In a context of auditory attention decoding, we find that a greedy channel selection based on the utility metric does not show a significant optimality gap compared to optimal channel selection, whereas other state-of-the-art greedy or *l*_1_-norm penalized methods do show a significant loss in performance. Furthermore, we demonstrate that the MIQP formulation also provides a natural way to incorporate topology constraints in the selection, e.g., for electrode placement in neuro-sensor networks with galvanic separation constraints. Furthermore, a combination of this utility-based greedy selection with an MIQP solver allows to perform a topology constrained electrode placement, even in large scale problems with more than 100 candidate positions.

## I. Introduction

Electroencephalography (EEG) is a popular non-invasive technology to record macro-scale electrophysiological activity in the brain. Most high-end EEG systems record from 20 up to 256 scalp electrodes [1], [2]. While using a large number of electrodes allows to record at a high spatial resolution, such high-density recordings also come with several disadvantages; they require more expensive equipment, they lead to longer set-up times, they require more data storage/processing, and the higher dimensionality may cause overfitting in data-driven algorithms. Furthermore, when making the transition towards wearable EEG applications with devices that measure EEG during daily-life activities [3]–[8] [9], [10], a low channel count is important for miniaturization and to minimize power and bandwidth requirements [10], [11]. Therefore, there is a need for efficient and robust data-driven channel selection or electrode placement methods to reduce the number of EEG channels while having minimal impact on the application performance. In this paper, we focus on the channel selection problem for least-squares (LS) based neural decoding. For illustrative purposes, we tackle and analyze the channel selection problem in the context of speech decoding, in particular in an auditory attention decoding (AAD) task [1]. However, we keep the methodology sufficiently generic, thereby making it applicable to any LS-based neural decoding task.

EEG channel selection is a combinatorial problem of which the complexity increases exponentially with the number of channels, thereby making an exhaustive search over all possible channel combinations infeasible. For example, finding the best combination of 8 channels from a pool of 64 EEG channels requires evaluating more than 4 × 10^9^ possible combinations. If one can evaluate^1^ a single combination in 0.01 second, it would take 1.4 years to go over all combinations. That is why channel selection is typically tackled by suboptimal heuristic methods that can be computed in a realistic time frame.

EEG channel selection methods are broadly classified into two categories, namely, filtering methods and wrapper methods [12]. Filtering methods rely on the use of distance, information or correlation measures independent of the problem’s objective function to select the best subset of EEG channels[12], [13]. Wrapper methods, on the other hand, tries to explicitly optimize the problem’s objective function while performing channel selection, which is why they typically perform better than filtering methods.

Therefore, in this paper, we focus on widely used wrapper methods for channel selection in LS-based neural decoding. Typical wrapper methods for LS decoding solve the channel selection problem with approximate convex relaxation techniques such as the least absolute shrinkage and selection operator (LASSO) [14], or heuristic techniques such as an iterative greedy elimination based on, e.g., decoder weights [15]–[17] or LS-based channel utility [9], [18]. In [9], the greedy channel selection using the LS-utility metric was found to perform the best in terms of AAD performance compared to other suboptimal strategies but the performance gap compared to the truly *optimal* channel selection, from hereon referred to as the optimality gap, remains unknown.

In this paper we propose a globally optimal channel selection method by reformulating the channel selection problem to a mixed-integer quadratic program (MIQP), thereby allowing it to be solved by state-of-the-art MIQP solvers to find the *exact* solution of the combinatorial problem. While the computation time for solving the MIQP is still very high (too high for practical use), it is at least practically feasible as opposed to a brute-force exhaustive search. The resulting optimal channel selection allows to quantify the optimality gap of the afore-mentioned sub-optimal techniques in a specific application. In the context of an AAD task, we demonstrate that, unlike (group)-LASSO or decoder weight-based greedy selection, the greedy method based on the LS-utility metric does not perform significantly worse than the optimal MIQP-based channel selection, while improving 3-4 orders of magnitude in computation time.

While channel selection can be done post-hoc to reduce the dimensionality of the data, it can also be used for electrode placement in a context of wearable EEG. Considerable research is ongoing to make wearable miniature-EEG (mini-EEG) devices which allow to record EEG 24/7 in daily-life activities [3]–[8]. Although these mini-EEG devices only cover small skin areas due to their far-driven miniaturization, the concept of neuro-sensor networks enables the simultaneous use of multiple such mini-EEG devices connected wirelessly thereby increasing the spatial resolution [10], [19]. Such a collection of wirelessly interconnected mini-EEG devices is also known as wireless EEG sensor networks (WESNs). In this case, it is essential to find the best scalp locations to place these mini-EEG devices (or ‘nodes’), where each node consists of at least two closely spaced electrodes to locally record EEG. In [9], [14], WESN nodes were emulated by first generating a highly redundant set of candidate nodes by re-referencing high-density cap-EEG electrodes with neighboring electrodes followed by node selection. However, no topological constraints were imposed during node selection, which may result in practically infeasible WESN topologies [9]. For example, since the nodes correspond to physically separated mini-EEG devices, which are galvanically separated from each other (not connected by a wire), the selected WESN nodes are not allowed to share electrode locations. Hence, there is a need to include such constraints in channel and node selection methodologies and explore their impact on neural decoding performance. To this end, we show how such topological constraints can be incorporated in the aforementioned MIQP formulation. This also allows to analyze the impact of such a galvanic separation by solving the MIQP with and without such constraints.

Although the proposed MIQP-based optimal selection method allows to solve the full combinatorial problem in a feasible time, it again becomes practically infeasible when the number of channels/nodes to be selected is large (> 10) or if the total number of candidate channels/nodes is large (> 100). Therefore, we also propose a hybrid method to perform node selection by combining the greedy utility-based channel selection with an MIQP solver, where the former initially reduces the candidate set of channels/nodes to a smaller set that can then be processed by an MIQP solver in a reasonable amount of time. The resulting combination yields a practical method for channel/node selection with topology constraints. For the case of AAD, we show that the inclusion of this greedy ‘preprocessing’ does not create a significant optimality gap compared to a globally optimal selection.

The outline of the paper is as follows. In Section II, we review the the channel selection problem for LS-based neural decoding, and reformulate it as a (constrained) MIQP. In Section III we describe the experimental setup used in this work, and the performance evaluation strategy used to compare different channel selection methods. In Section IV, we report the results of the comparative analysis in terms of AAD performance and computation time. We discuss the results in Section V and we draw conclusions in Section VI.

### Note on terminology

The following terminology will be used consistently in the remainder of this paper. A *channel* is an EEG signal that originates from a single electrode pair over which the scalp potential is measured. A *node* represents a group of (at least two) closely spaced EEG electrodes such as those included in a wireless mini-EEG sensor device, emulated here as a group of nearby cap-EEG electrodes. In all our experiments reported in this paper, we only consider single-channel nodes consisting of a single electrode pair although all results can be extended to multi-channel nodes with more than 2 electrodes [9].

## II. Channel or Node selection for neural decoding

### A. Least-squares based Neural Decoding

Several studies have established that the neural responses of a subject measured as multi-channel EEG can be decoded to reconstruct certain features of the stimulus. For speech decoding in particular, it has been found that least-squares based linear regression models allow to reconstruct different representations of the auditory speech stimulus such as the speech envelope [1], [2], [15], the spectrogram, phonetic features [20], etc. Moreover, neural decoding of EEG to speech envelopes has been used in auditory attention decoding (AAD) algorithms which allow to determine to which speaker a subject is attending when listening to a mixture of speakers in a so-called cocktail party scenario [1], [2], [15].

The neural decoding problem consists of finding a spatio-temporal decoder 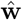 which linearly combines the EEG data to reconstruct the stimulus **d** in least squares (LS) sense:

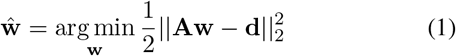

where **A** is a *T × QC* matrix containing *T* time samples of the *C* EEG channels and *Q* − 1 non-causal time-lagged copies of each channel in its columns. Each channel and its *Q -* 1 time-lagged copies are assumed to be grouped in adjacent columns in the matrix **A**. The time-lagged copies are added to cope with time delays and convolutive responses [1], [2], [15]. The problems where time-lagged copies are not used can be considered as a special case of (1) where *Q* = 1.

In addition, in (1), **d** is a *T*-dimensional vector containing *T* time samples of a relevant representation of the stimulus which in the case of speech can be represented by the speech envelope, the spectrogram or even low level representations like phonemes [20]. In the present work, we address the problem of decoding the speech envelope.

The solution of (1) is given by

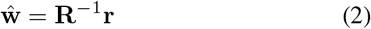

where **R** = **A**^*T*^ **A** and **r** = **A**^*T*^ **d**. If required, a diagonal loading term can be added to **R** as a regularization mechanism [1], although it was shown in [2] that this is not necessary (and to be avoided) in case sufficient training data is available to populate **R**.

The problem of selecting the best *N* (< *C*) channels which minimizes (1) is a combinatorial problem. Different approximate approaches have been proposed to solve this problem, for example, convex relaxations like (group-)LASSO[14] or greedy methods with iterative elimination of channels [9], [15]–[17].

### B. Greedy channel selection

Assuming the *Q* time-lagged copies of each EEG channel are in adjacent columns in the matrix **A**, we can define the following partitioning for the spatio-temporal decoder 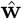:

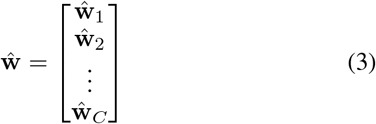

with the subvectors 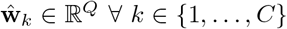, the decoder coefficients corresponding to *k*-th channel and its *Q* copies. In [15]–[17], the EEG channels are iteratively removed one by one in a greedy fashion by every time deleting the channel *k* for which the *l*_2_-norm 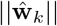 is the smallest. After each iteration, the optimal decoder is recomputed based on the remaining channels until the desired number of *N* channels is reached. In the remaining of this paper, we will refer to this method as the decoder magnitude-based (DMB) greedy method or DMB-G.

However, in [21] it was argued that the magnitude of the entries in the decoder 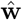 do not necessarily reflect the importance of the corresponding channel as it is scaling dependent and it does not properly take interactions across channels into account. Instead, it was argued to quantify the importance or ‘utility’ of a channel *k* by the increase in the least squared error (LSE) if channel *k* were to be removed and the decoder would be fully re-optimized. In [21] an efficient computation for this utility metric was proposed, and it was shown in [9] that it outperforms the DMB metric in a greedy channel selection procedure. We will refer to this method as the utility-based (UB) greedy channel selection method^2^ or UB-G, which we will briefly review below as it will be part of the hybrid method proposed in Section II-D3.

Since the matrix **A** in (1) contains *Q* time-lagged copies of *C*-channel EEG in its columns, the removal of an EEG single channel corresponds to the removal of a group of *Q* columns from **A**. The utility of a group of columns of **A**, referred to as the *group-utility*, is defined as:s

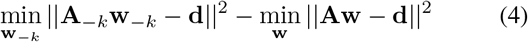

where **A**_*−k*_ is the matrix **A** with the *Q* columns corresponding to channel *k* removed. It has been shown that this group-utility, can be computed efficiently based on 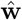 without having to compute the new optimal decoder 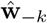 for each channel *k* [21]. In UB-G, this group-utility metric is used to iteratively eliminate channels with the least group-utility to select the ‘best’ *N* channels. To this end, assume without loss of generality (w.l.o.g.) that the channel *k* and its time-lagged copies for which we compute the group-utility corresponds to the last *Q* columns of **A**.

Defining the block partitioning of **R**^−1^ in (2) as:

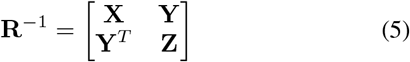

where **Z** is a *Q × Q* matrix corresponding to the *Q* time lags associated with channel *k* (here assumed to be in the *Q* last columns of **A** w.l.o.g.). The group-utility of channel *k* can be efficiently computed as [9], [21]:

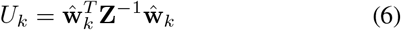

where 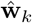 contains the last *Q* entries of 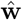. It can be shown that (6) leads to the exact same quantity as defined in (4) [21] without the need to recompute (2), which would involve a large matrix inversion for each candidate channel removal.

To select *N* (out of *C*) channels of EEG data used in the neural decoding problem (1), UB-G uses the algorithm illustrated in Fig. 1. First, the group-utility of each of the *C* channels is computed using (6) followed by the removal of the channel with the least group-utility. After this removal, 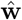 is recomputed using (2) but now with the (*C* − 1) channel EEG data. The new group-utilities of each channel in the new (*C* − 1) channel set are re-computed from (6), again followed by removal of the channel with the least group-utility. The procedure is repeated until only *N* channels remain.

**Fig. 1.**
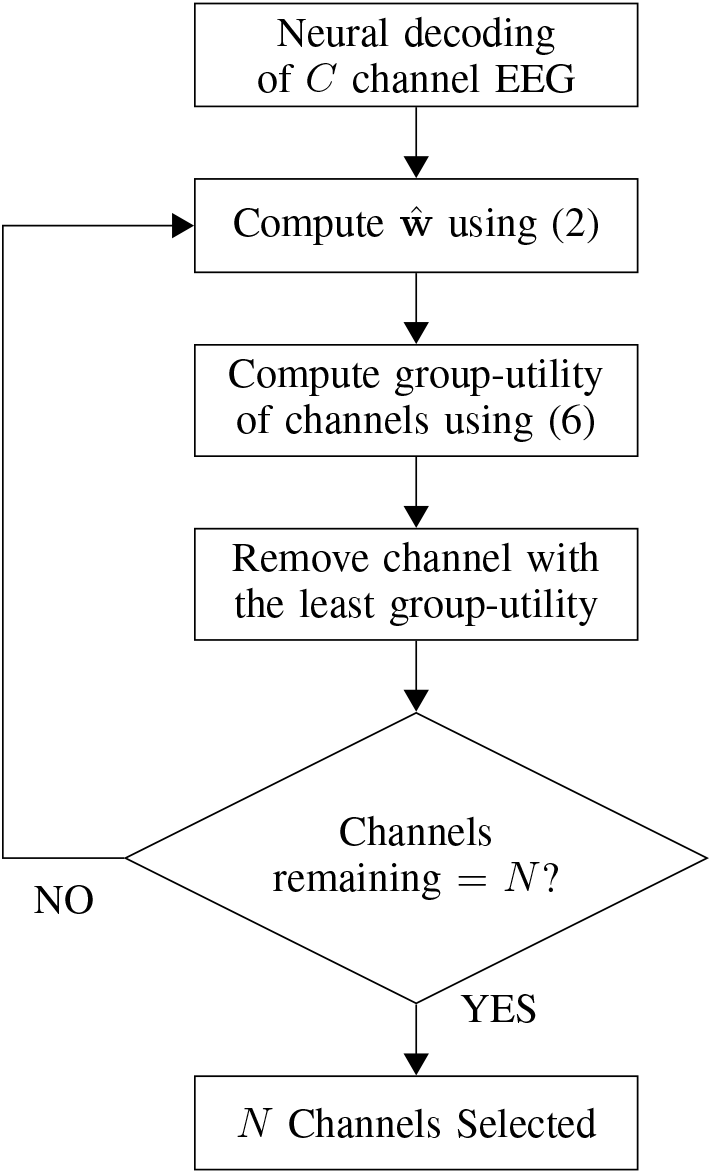
Utility-based greedy (UB-G) channel selection algorithm.

### C. Optimal channel selection

While UB-G outperforms other state-of-the-art channel selection methods [9], it still leads to sub-optimal solutions to the channel selection problem due to its greedy approach (as not all possible combinations of *N* channels are considered). As such, other combinations of *N* channels may lead to even lower squared errors. The optimality gap between UB-G channel selection and the optimal selection that truly minimizes the LS cost remains unknown since it requires investigating all possible channel combinations, which is computationally infeasible. In this section, we propose a feasible method to find this global optimum of the channel selection problem.

We introduce a boolean vector **z** of length *C* defined as **z** = [*z*_1_ *…, z_C_*]^*T*^ with *z_k_* ∈ {0, 1} ∀ *k* ∈ {1*, …, C*}, which contains selection variables for each channel. Now, modifying the optimization problem in (1) with the newly introduced variable **z**, the channel selection problem can be equivalently formulated as:

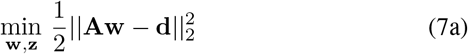

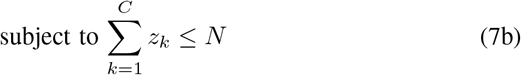

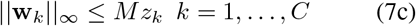

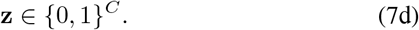

where **w**_*k*_ is the subvector defined in (3), *M* is a sufficiently large ^3^ positive integer and || · || _*∞*_ is the *l*_*∞*_ norm.^4^

The constraint in (7b) ensures that at most *N* entries of **z** assumes the value of 1, thereby selecting *N* channels. The constraint (7c) imposes that the entries of **z** act like selection variables for the columns of **A**. A value of 0 at *z_k_* forces all entries of **w**_*k*_ to be 0, thereby removing the *Q* columns of **A** corresponding to channel *k* from the problem. On the other hand, a value of 1 at *z_k_* gives the freedom for all entries of **w**_*k*_ to assume non-zero values, thereby selecting *Q* columns of **A** corresponding to channel *k*.

The optimization problem of the kind in (7a)–(7d) is an instance of a so-called mixed-integer quadratic program (MIQP). With the help of solvers like CPLEX[22], Gurobi [23], etc. the MIQP (7a)–(7d) can be solved to optimality for moderate values of *C* (< 100) and for small values of *N* (< 10), in feasible time. In the experiments in this paper, we used the Gurobi solver [23] to solve (7a)–(7d). From this solution, we considered the channels which correspond to non-zero entries of **z** as the optimal channels. We will refer to this optimal channel selection method as OCS in the sequel.

### D. Node placement with galvanic separation constraints

In neuro-sensor networks such as WESNs, the selection of EEG channels usually has to satisfy certain topological constraints. For example, the *N* nodes of a WESN correspond to stand-alone mini-EEG sensors which are not connected by a wire, i.e., they are galvanically separated. This means that the *N* selected nodes are not allowed to share the same electrodes. In this subsection, we describe how such constraints can be included in the OCS method.

#### 1) WESN Emulation

We use the procedure of [9] to emulate candidate WESN nodes from a 64-channel standard cap EEG recording. Without loss of generality, we only address the case of a WESN made up of single-channel nodes, i.e., each node consists of two electrodes separated by a short distance. These nodes are selected from a set of candidate nodes created by pairing each electrode of the *C*-electrode cap with each of its nearby electrodes that are at a distance of at most *r* cm, where *r* is the desired maximum span between the electrodes within a single WESN node. Using this criteria, a set of *P* candidate single-channel node locations and orientations were generated from the original *C* electrodes. Since each node then corresponds to a single electrode-pair, it contributes a single channel of EEG. Hence, the node selection or node placement problem while constructing a WESN can be viewed as a *channel selection* problem as described in Section II-A, where *C* is replaced by *P* (note that in practice *P* ≫ *C*, which means the set of candidate nodes is highly redundant).

#### 2) OCS method for galvanically separated node selection(OCS-GS)

The OCS method described in Section II-C can be used to find optimal node locations for WESNs by selecting *N* nodes from *P* candidate nodes. The problem formulation (7a)–(7d) can perform optimal node selection by replacing **A** with **A**_*P*_ where now **A**_*P*_ contains *Q* time-delayed copies of EEG signals from each of the *P* nodes in its columns. Assume *ε* ⊂ {1, …, *P*} × {1, …, *P*} is the set that contains all ordered pairs of nodes sharing an electrode, i.e., all node pairs that are galvanically connected. To select galvanically separated nodes, the problem given in (7a)–(7d) is modified as below:

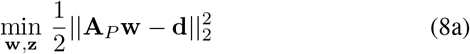

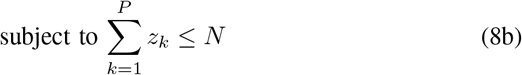

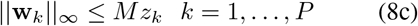

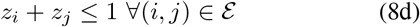

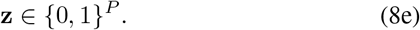

Here, the constraint (8d) ensures that if node *i* and *j* share an electrode, only one of them will be selected. The optimization problem (8a)–(8e) is again an MIQP, which can be solved with the Gurobi solver [23]. We will refer to this modified optimal node selection method with galvanic separation constraint as OCS-GS.

#### 3) Hybrid method for galvanically separated node selection

In the previous section, we described an optimal node selection method for optimal node placement with galvanic separation constraints. However, the computational time required to obtain these optimal node selections is generally too high for practical purposes due to large values of *P* (compared to *C*). In this section, we propose a hybrid node selection strategy involving greedy components to perform node selection with galvanic separation constraints with more reasonable computation times.

Note that the set of *P* candidate nodes is a redundant set, which will make the matrix **R** in (2) rank deficient, thereby hampering the computation of (2)–(6) in the DMB-G and UB-G method. For the case of DMB-G, a pseudo-inverse can be used in (2). For the UB-G method, an extension of the utility metric for such rank-deficient problems is proposed in [21] based on a minimum-norm criterion, which was also used in [9] for the node selection problem. We will use a similar fix when we apply UB-G, and we refer to [9] or [21] for further details. The gLASSO and the MIQP solvers do not explicitly compute **R** or its inverse, and therefore do not require a fix when the latter is singular.

We first apply UB-G to prune the *P* candidate nodes to *K* nodes with *N* ≪ *K* < *P*, where *N* is the number of nodes to be selected and where *K* is a value which is small enough such that OCS or OCS-GS can be computed in a reasonable amount of time. The pruning stage is followed by node selection from the *K* remaining candidate nodes with galvanic separation constraints using the OCS-GS method in (8a)–(8e). The pruning stage is applied to reduce the number of candidates on which OCS or OCS-GS is computed. Meanwhile, the use of OCS-GS for node selection ensures that the solution is near-optimal while the galvanic separation constraints are satisfied. We refer to this method as ‘hybrid’ in the remaining of the paper.

## III. Experimental set-up

### A. Description of EEG dataset

The data set used for the experiments reported in this paper, originally described in [2], The data used for the analyses in this paper consist of 64-channel EEG recorded using a BioSemi ActiveTwo system from 16 subjects who sat through three experiments within a single recording session. During each experiment, the subjects listened to two simultaneous children stories narrated by two different male speakers coming from two distinct spatial locations (left and right of the subject), and were asked to attend to only one of them while ignoring the other. The first two experiments each included four presentations of different six-minute story parts (the unattended speaker from the first experiment becomes the attended speaker in the second experiment and vice versa). This results in 2 × 4 × 6 = 48 minutes of EEG data. The third experiment consisted of four shorter presentations of the first two-minutes of the same four story parts. These presentations were repeated three times, to build a set of recordings of repetitions, thereby adding 24 extra minutes of EEG data to obtain 72 minutes of EEG data in total per subject. A version of this dataset is available online along with a more detailed description [24]. During preprocessing we re-referenced the EEG data to the Cz electrode.

In Section II-D1, we briefly described the procedure for WESN emulation, originally used in [9]. We created the candidate two-electrode single-channel nodes for WESNs with a maximum distance of *r* = 5*cm* between the electrodes. This corresponds to the configuration used in [9] ensuring that a large number of candidate node locations and orientations are generated but at the same time the electrodes of each pair have a reasonably short distance between them to emulate a miniaturized EEG-sensor node. This resulted in *P* = 209 candidate nodes with an average inter-electrode distance of 3.7 cm.

### B. AAD performance evaluation

For validation of the different channel selection algorithms described in this paper, we used the AAD procedure from [2]. First, we estimated a subject-dependent linear spatio-temporal decoder 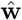 based on (1) where **d** contains the envelope of the attended speaker. We filtered both the EEG data and the speech envelope using a bandpass filter between 1 − 9Hz and we followed it by downsampling both to 20Hz. In the comparison between UB-G and OCS on the standard cap-EEG channel selection, we used the value of *Q* = 6, which corresponds to time delays up to 250*ms* for both channel selection as well as performance evaluation. It has been shown that the time delays up to 250*ms* are the most effective for reconstructing envelopes using EEG for the sake of attention decoding [1], [2]. Within these delays, the delays between 140*ms* and 200*ms* have been shown to be the most discriminative to decode auditory attention to speech [1], [25]. We selected the lower value of *Q* in this case to solve the MIQP-based optimal node selection in feasible time, as reducing *Q* leads to a lower number of total variables in the node selection problem. For the hybrid method, a larger value of *Q* is possible, but we also set it to *Q* = 2 in order to compare and quantify the potential optimality gap with OCS and OCS-GS.

We use a leave-one-trial-out cross-validation scheme, in which the data is split in *L* trials of 60*s*. We used each trial once as a test trial. When testing on trial *l*, we compute (1) on the entire EEG recording after cutting out trial *l* from **A** and **d**, to find the decoder 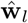 for test trial *l*. The decoder 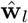 is used to reconstruct the attended speech envelope for trial *l* using:

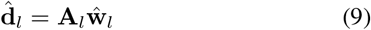

Once the attended speech envelope has been estimated, we found the Pearson correlation coefficients between 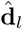 and the attended and unattended speech envelopes in trial *l* as *r*_*a*_ and *r_u_* respectively. We considered a trial to be successfully decoded if *r_a_ > r_u_*. We used the percentage of successfully decoded trials as the AAD performance measure (analyzed per subject). Similar to [18], we also report the mean attended correlation coefficients *r_a_* (averaged across trials within a subject). The mean attended correlation is larger when the reconstruction in (1) is better.

## IV. Results

### A. AAD Performance Analysis

#### 1) Channel selection in standard cap EEG

We compared the optimal channel selection (OCS) method to three different approximate EEG channel selection strategies for least-squares based neural decoding, namely group-LASSO (gLASSO)^5^ [14], and a greedy selection based on either decoder magnitude (DMB-G) [15], [16] or group-utility (UB-G) [9], [21], to select *N* = 1, 2, …, 8 standard cap-EEG channels in an AAD task. The decoding accuracy and mean attended correlation across subjects are plotted in Fig. 2a and Fig. 2b respectively. Note that running OCS for values of *N* ≥ 10 takes an unacceptably long time, which is why we exclude these cases from the analysis. However, it is noted that the performance of the OCS-based selected channels for *N* = 8 is close to the performance with all 63 channels, so the OCS is expected to reach the full-channel performance for *N* ≥ 10. We compared each of the approximate methods to OCS using linear mixed-effects (LME) models with the number of channels (*N*) and the two methods (OCS and an approximate method) as fixed effects, and subjects as random effect. We used the software R (version 3.6.2), and the R package ‘nlme version 3.1-144’ [27] for fitting the linear mixed effect models. All the linear mixed effect models in this work were fitted by maximizing the restricted log-likelihood, and the residuals were checked for normality to ensure a good fit. When we compared DMB-G and gLASSO to OCS, we found these approximate methods to be significantly different from OCS with *p*–values < 0.001 for both decoding accuracy and attended correlation comparisons. However, when we compared UB-G to OCS there was no significant difference in decoding accuracies (*p* = 0.63) and mean attended correlation (*p* = 0.16) between the two methods. In Fig. 3, the distribution of the best Cz-ref channels selected by the OCS and UB-G method across 16 subjects are shown in the form of a heatmap topoplot.

**Fig. 2.**
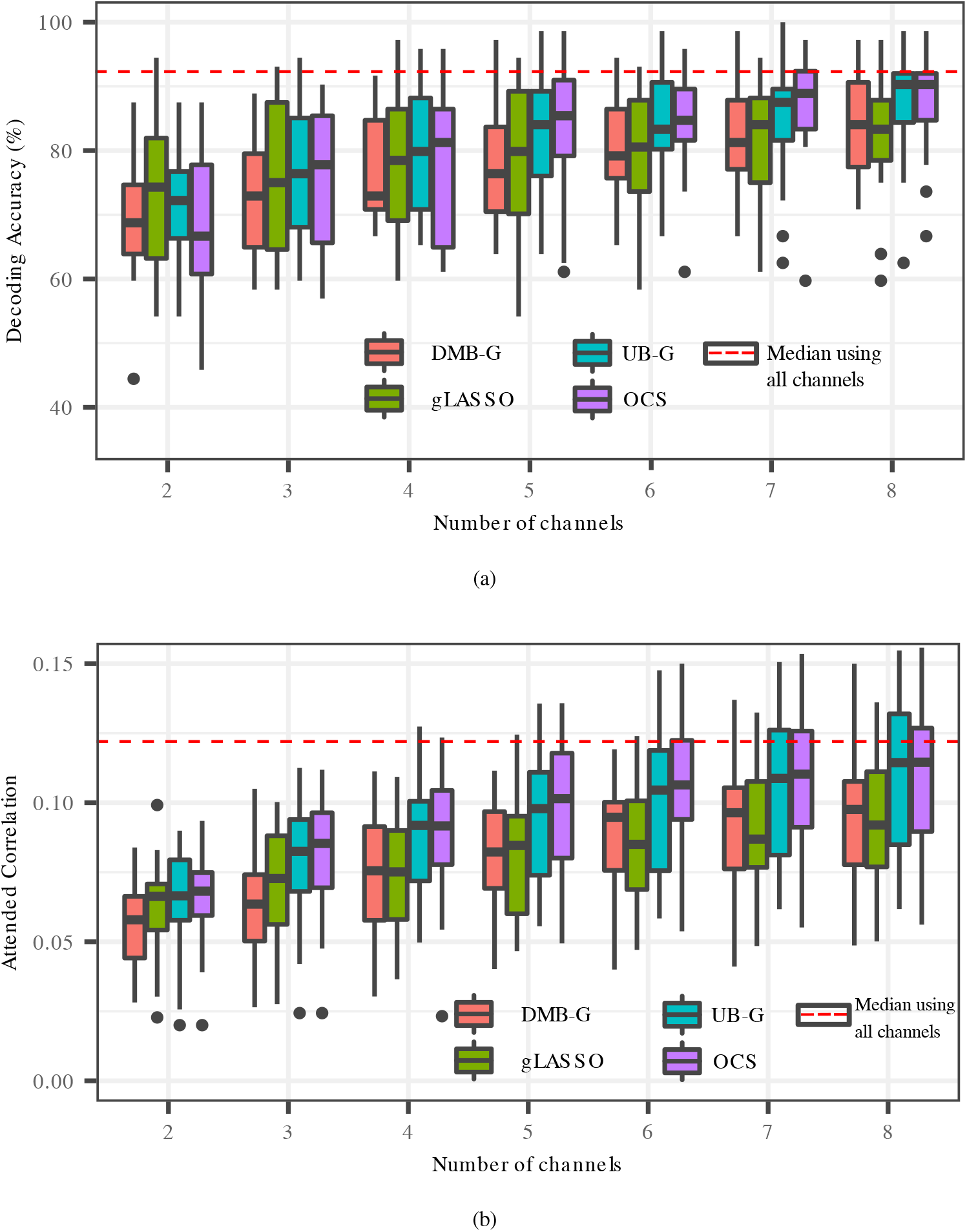
OCS compared to all the approximate methods, namely gLASSO, DMB-G and UB-G, with respect to (a) the decoding accuracy (b) the mean attended correlation. The black dots represent outliers, values beyond 1.5 × *IQR* (inter quartile range) from the quartiles.

**Fig. 3.**
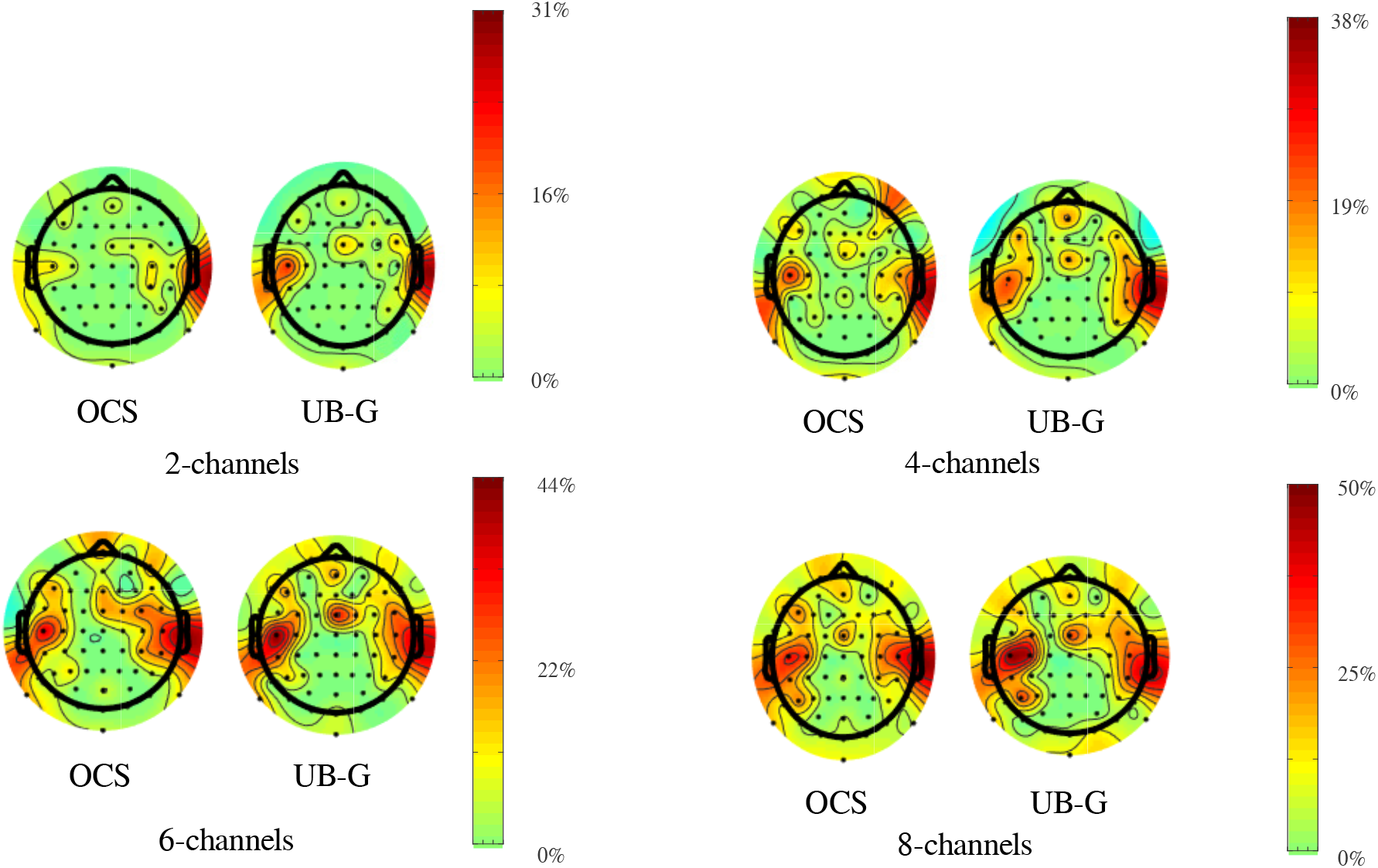
Heatmap topoplots illustrating the scalp locations of the best Cz-ref channels selected by the OCS and UB-G methods across all subjects. The color bar indicates the percentage of 16 subjects selecting an electrode.

#### 2) Node selection in WESNs with galvanic separation constraints

Next, we used OCS and its modified version of OCS-GS in (8a)–(8e) to investigate the impact of galvanic separation of nodes of a WESN on AAD performance. Fig. 4 demonstrates this comparison. Please note that here the comparison is not between two methods (in both cases we performed an optimal selection based on an MIQP) but between the two scenarios, namely node selection with galvanic separation (GS) constraints and without galvanic separation (NGS) constraints. Due to the size of the MIQP with *P* = 209 candidate channels, we could only find solutions in a feasible computation time for *N* ≤ 6 selected channels and for *Q* = 2 (see also Section III-B). For *N* = 5, we had to exclude four subjects and for *N* = 6, we had to exclude five subjects, as the solver could not find an optimal solution for these subjects due to numerical issues. For all the other cases of *N*, all 16 subjects have been included in the comparisons.

**Fig. 4.**
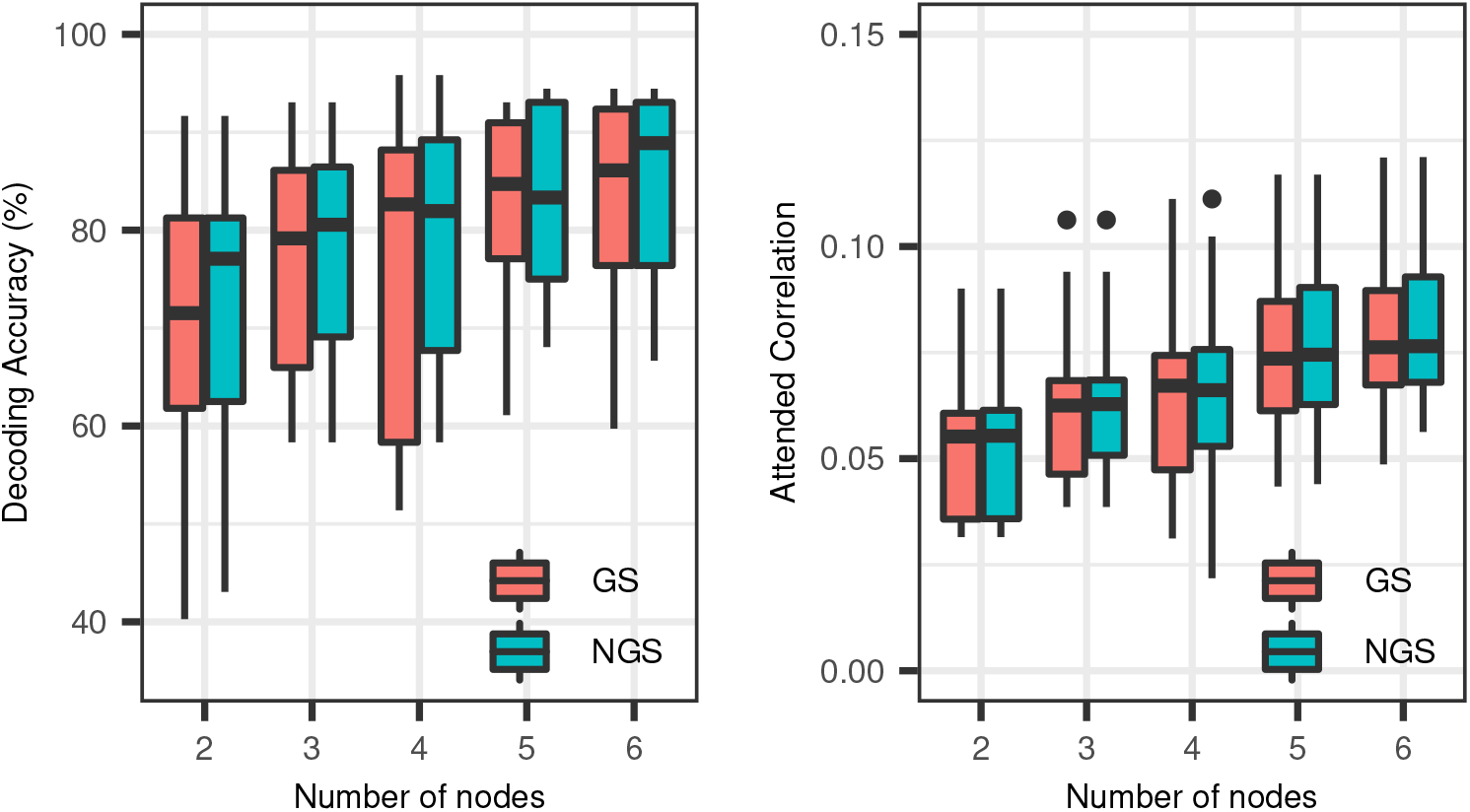
Optimal WESN node selection with galvanic separation (GS) and without galvanic separation (NGS) constraints: OCS-GS as in (8a)–(8e) was used in the former and OCS as in (7a)–(7d) was used in the latter. In both cases *Q* = 2 with sample delays 150*ms* and 200*ms*. The black dots represent outliers, values beyond 1.5 × *IQR* from the quartiles.

Fig. 4 demonstrates that the galvanic isolation between nodes of a WESN has no negative impact on AAD performance, which we also confirmed using statistical tests. We used Wilcoxon signed rank tests^6^ with and without Holm-Bonferroni correction to compare the decoding accuracies and attended correlation for each value of *N*. We observed no significant differences between the two scenarios with respect to decoding accuracies and attended correlation. The Wilcoxon signed rank test *p*–values and Holm-Bonferroni corrected *p*-values are provided in Table I.

**TABLE I.** *p*–values of pairwise comparisons of AAD performances, decoding accuracy (Dec.Acc) and attended correlation (Att. Corr.), using a Wilcoxon signed rank test with and without Holm-Bonferroni (HB) correction.

| Comparison | $N$ | Dec. Acc. | | Att. Corr. | |
| --- | --- | --- | --- | --- | --- |
|  |  | no HB | HB | no HB | HB |
| GS<br>vs<br>NGS | 2 | 0.06 | 0.35 | 0.43 | 1 |
|  | 3 | 0.35 | 1 | 0.37 | 1 |
|  | 4 | 0.23 | 1 | 0.54 | 1 |
|  | 5 | 0.81 | 1 | 0.21 | 1 |
|  | 6 | 0.21 | 1 | 0.03 | 0.2 |
| OCS-GS<br>vs<br>Hybrid | 2 | 0.70 | 1 | 0.89 | 1 |
|  | 3 | 0.98 | 1 | 0.15 | 1 |
|  | 4 | 0.33 | 1 | 0.52 | 1 |
|  | 5 | 0.57 | 1 | 0.45 | 1 |
|  | 6 | 0.61 | 1 | 0.96 | 1 |
| Hybrid<br>vs<br>UB-G | 2 | 0.4 | 1 | 0.03 | 0.2 |
|  | 3 | 0.82 | 1 | 0.30 | 1 |
|  | 4 | 0.99 | 1 | 0.46 | 1 |
|  | 5 | 0.91 | 1 | 0.87 | 1 |
|  | 6 | 1.00 | 1 | 0.60 | 1 |
|  | 7 | 0.83 | 1 | 0.60 | 1 |
|  | 8 | 0.59 | 1 | 0.06 | 0.37 |

We used the hybrid method proposed in Section II-D3 to perform node selection with galvanic separation constraints. The pruning stage of the hybrid method pruned *P* = 209 candidate nodes to *K* = 64 candidate nodes. We applied the OCS-GS method to select *N* nodes from this set of *K* candidate nodes with galvanic separation constraints. In order to evaluate a possible optimality gap between the hybrid and the OCS-GS methods, we compared the AAD performance and the mean attended correlation across subjects between both methods for WESN node selection with galvanic separation constraints. The results are shown in Fig. 5, which suggest that the hybrid method performs very similar to the optimal methods, both in terms of decoding accuracies (Fig. 5a) and mean attended correlation (Fig. 5b). A Wilcoxon signed rank test, with and without Holm-Bonferroni corrections, failed to reject the null hypothesis when comparing both node selection methods with galvanic separation constraints. The *p* values can be found in Table I.

**Fig. 5.**
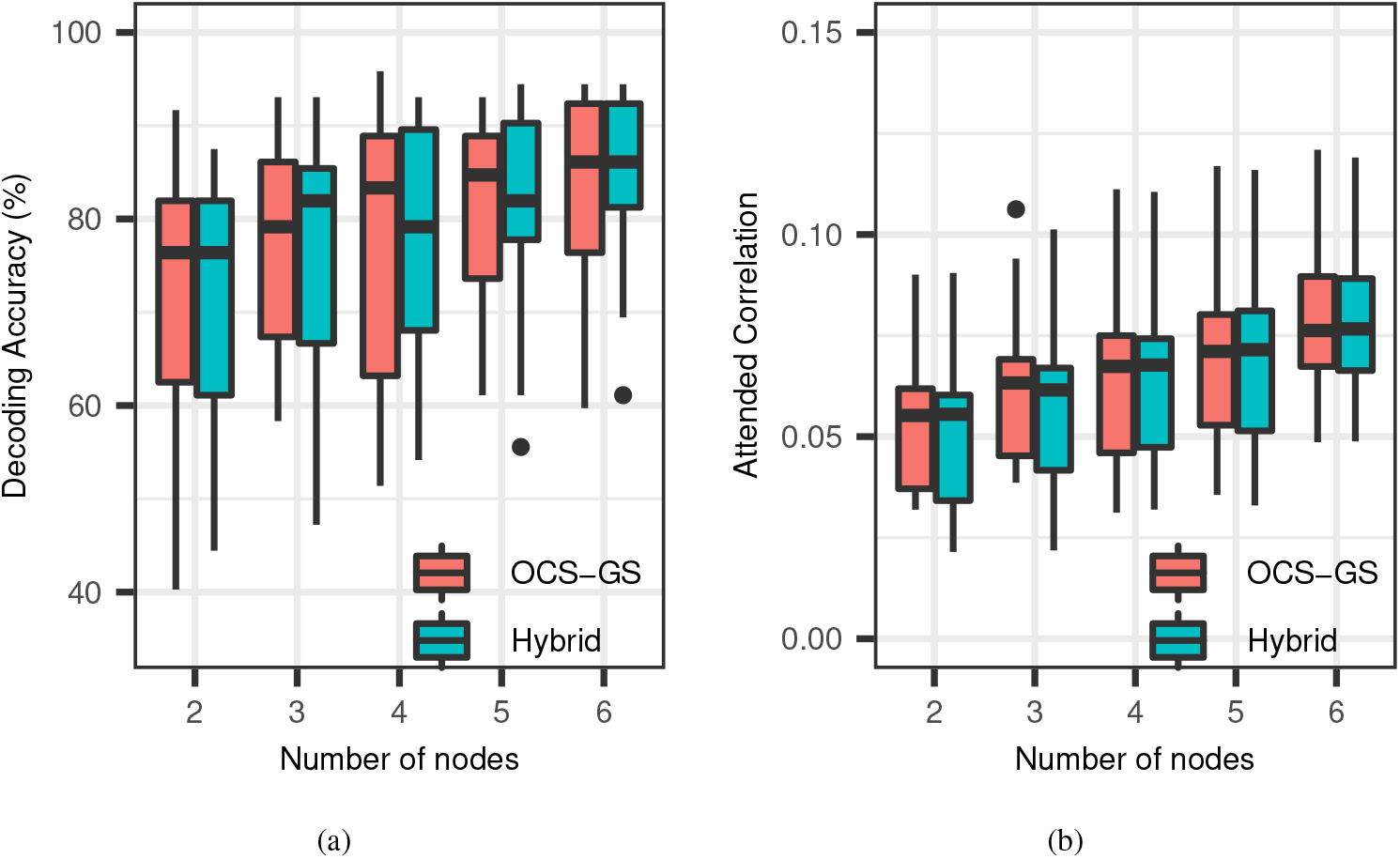
Galvanically separated node selection: Optimal node selection with galvanic separation constraints (OCS-GS) compared to hybrid node selection with galvanic separation constraints. *Q* = 2 with sample delays 150*ms* and 200*ms*. The black dots represent outliers, values beyond 1.5 × *IQR*, from the quartiles.

Since we now established that the hybrid method performs equally well as the OCS-GS method, we can use the hybrid method to re-investigate the impact of galvanic separation of nodes. The hybrid method now allows to select more channels (up to *N* = 8) and to include all sample delays in the decoder up to 250*ms* (*Q* = 6). The results are shown in Fig. 6. The figures seem to indicate little to no effect of galvanic separation on AAD performance, which confirms the earlier analysis in Fig. 5 for *Q* = 2 and *N* ≤ 6. A Wilcoxon signed rank test without Holm-Bonferroni corrections, failed to reject the null hypothesis in all but one case of *N* = 2 for attended correlation (*p* = 0.03). However, with Holm-Bonferroni correction, the Wilcoxon signed rank test did not show a significant difference for this case. All the *p* values can be found in Table I.

**Fig. 6.**
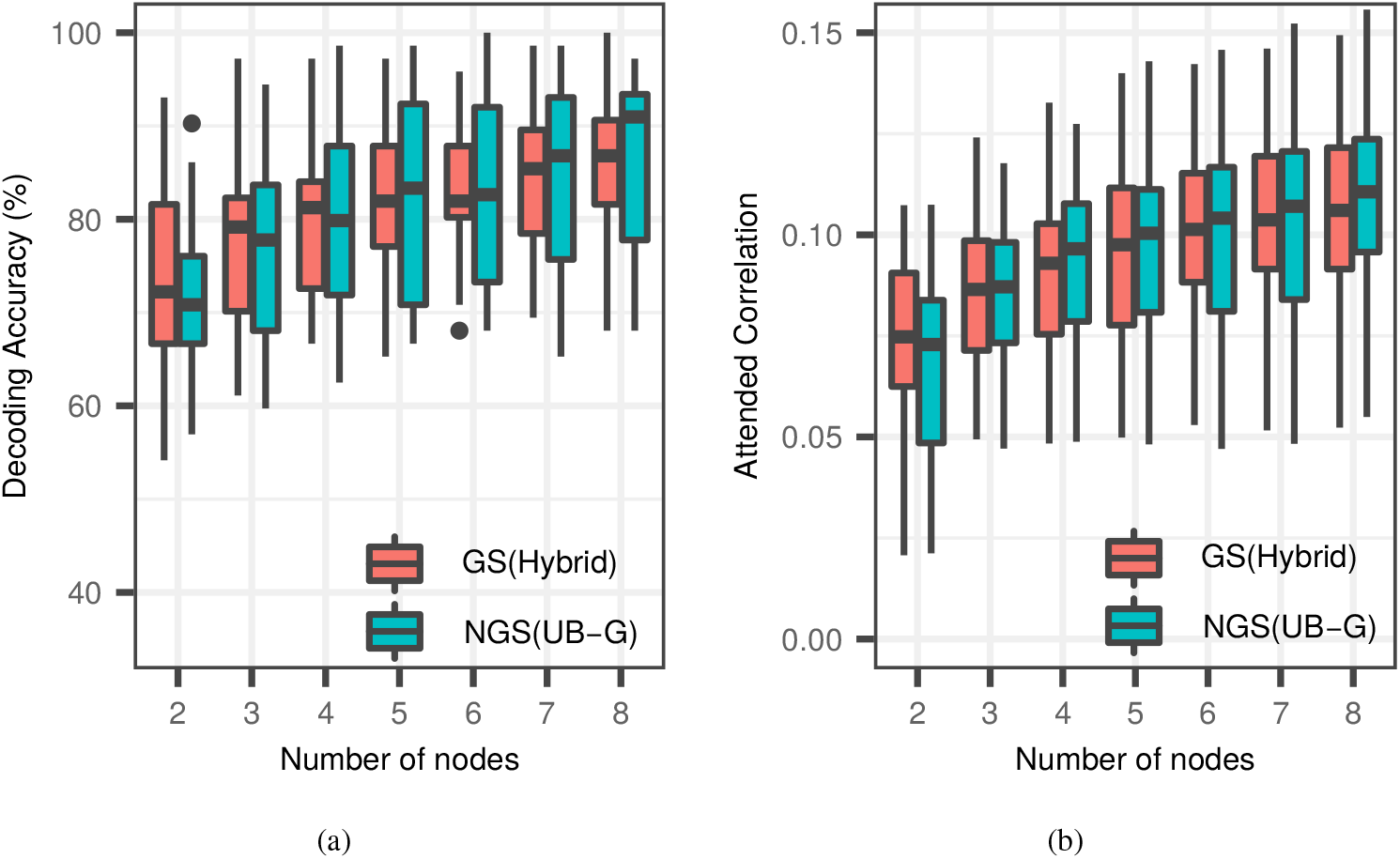
Galvanically separated node selection: Hybrid node selection with galvanic separation constraints compared to UB-G node selection without galvanic separation constraints. *Q* = 6 with sample delays 0*ms* to 250*ms*. The black dots represent outliers, values beyond 1.5 × *IQR*, from the quartiles.

### B. Computation Time Analysis

In Fig. 7, we show a comparison of the computation times to select *N* = 1, 2 …, 6 WESN nodes(out of *P* = 209) without galvanic separation using OCS, Hybrid (where the OCS-GS step is replaced with an OCS step for fairness) and UB-G. The computations were carried out using MATLAB R2018b on an Intel^®^ Xeon^®^ CPU clocked at 2.50GHz. The expected computation time when performing an exhaustive search over all possible channel combinations for optimal selection is also plotted. A computation time of 0.01 seconds was assumed for evaluating one combination, which is the actual time taken on an Intel^®^ Xeon^®^ CPU clocked at 2.50GHz using MATLAB R2018b. Compared to the exhaustive search, the OCS method clearly finds optimal solutions in computationally feasible time for small values of *N*. Nevertheless, the computation time of OCS, and the hybrid method increases exponentially with *N* (linearly on a logarithmic scale as in Fig. 7). However, for *N* > 3, the hybrid method’s computation time is smaller than OCS by at least an order of magnitude. The hybrid method prunes the *P* = 209 candidate nodes *P* by to a set of only *K* = 64 before applying OCS. In addition, we can clearly observe the advantage of UB-G over the other two methods. UB-G is much faster and its computation time does not increase with *N* due to the greedy implementation.

**Fig. 7.**
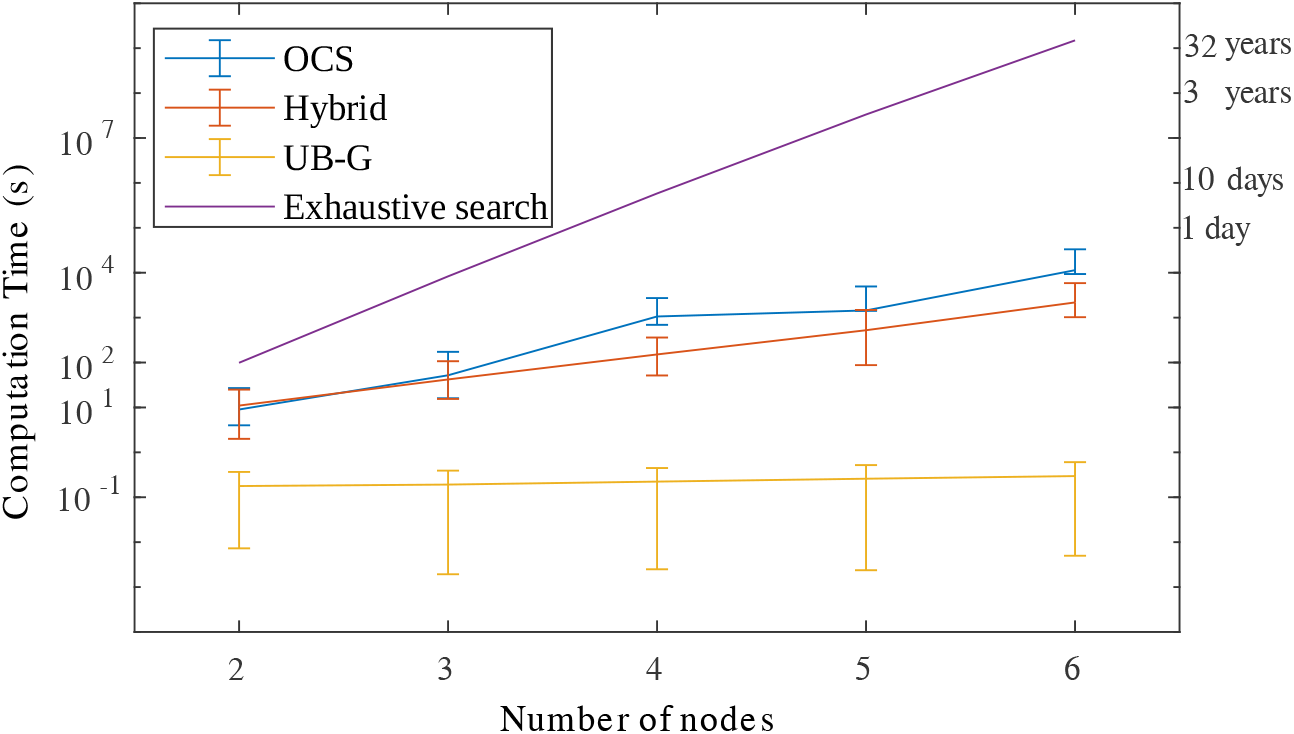
Log-plot of computation time comparison: OCS, Hybrid and UB-G, were used to select *N* = 1, 2, …, 6 WESN nodes without galvanic separation. The error bars denote the 25 – 75 percentile across subjects. The expected computation time when an exhaustive search over all possible channel combinations is also plotted. The y-axis labels on the right provide a perspective for the computation times of exhaustive search.

## V. Discussion

The first goal of this paper was to find exact solutions to the EEG channel selection problem for least-squares based neural decoding in a feasible time (as opposed to an exhaustive search over all possible combinations) thereby allowing us to quantify the potential optimality gap of three state-of-the-art approximate channel selection methods. To this end, we proposed an OCS method based on an MIQP formulation of the channel selection problem of which the solution is guaranteed to be the truly optimal selection of channels in least squares sense. Fig. 2a and Fig. 2b suggest that, among the approximate methods, UB-G is the only one which does not have a clear optimality gap with the OCS. Moreover, statistical tests seem to indicate no significant difference between the UB-G and OCS methods as detailed in Section IV whereas both the other approximate methods, namely DMB-G [15]–[17] and gLASSO [14], perform significantly worse than OCS. We note that this result of statistical testing only implies lack of sufficient evidence to reject the null-hypothesis, i.e., it does not guarantee the null-hypothesis to be true. Nevertheless, the large *p* values suggest that -in case there would be a optimality gap-it is at least very small compared to the natural spread across different subjects. In addition, the distribution of the electrodes selected by the OCS and UB-G methods, as shown in Fig. 3, also indicate that both methods tend to select electrodes from similar scalp locations in the majority of the subjects.

The results shown in Fig. 2a and Fig. 2b confirm the results in [9], where the advantage of UB-G over other approximate methods was already observed, yet a comparison with the optimal channel combination was not possible due to the infeasible computation time to test all possible channel combinations. Due to the MIQP formulation proposed in Section II, we were able to circumvent that problem, at least for values up to *N* = 8. As a reference, assuming 0.01 seconds for evaluating one combination, an exhaustive search over all channel combinations to find the best 8 channels from 64 would require 1.4 years. The lack of a significant optimality gap adds further support for using greedy selection using the LS-utility metric as a proxy for optimal selection. Furthermore, due to the algebraic trick provided in [21] resulting in the expression (6), the computation of the LS-utility metric is sufficiently cheap to be used in practice, as illustrated in Fig. 7.

While our initial analysis focused on channel selection in standard cap EEG, we also investigated the node selection problem in WESNs. WESNs are envisaged to use a multitude of mini-EEG devices thereby increasing spatial resolution and scalp coverage with full flexibility due to the absence of wires between the EEG sensor devices. Due to this absence of wires, the individual EEG sensor devices are supposed to be galvanically separated. To find the ideal locations for such individual mini-EEG devices in a WESN context and to study the impact of galvanic separation, the second goal of the paper was to perform node selection with the inclusion of topological constraints. In [14], the inclusion of topological constraints was explored but in an approximate group-LASSO framework using a heuristic. In Section II-D we modified the OCS method to include topological constraints in the LS optimization problem itself, again in the form of an MIQP. We used this approach, referred to as OCS-GS, to select an optimal set of *N* galvanically separated nodes to form a WESN. Furthermore, the impact of galvanic separation in WESNs was analyzed and in Fig. 4 and Fig. 6 it is shown that WESNs using galvanically separated nodes and WESNs constructed using nodes without galvanic separation constraint perform similarly. The statistical analysis reported in Table I also demonstrates this. These results are promising and reassuring to further investigate the use of WESNs for AAD, as they show that the absence of wires across the EEG sensors, thereby effectively decoupling their EEG content, does not affect the decoding performance when fusing the EEG activity across the different sensors.

However, a major problem of the topology-constrained MIQP in (8a)–(8e) is that it requires long computation times. As observed in Section IV-B, the optimal selection requires hours to complete for *N* > 3, with the time requirements increasing exponentially with the number of channels to be selected. On the other hand, the faster UB-G method does not allow to take topological constraints into account. Thus, we proposed a hybrid approach in Section II-D3 which utilizes the best features of both greedy (UB-G) and optimal (OCS) channel selection methods to result in a compromise in terms of computation time and optimality while also allowing to include topological constraints as in the OCS-GS method. We used the hybrid technique to perform node selection with galvanic separation constraints and this was compared with OCS-GS. The results shown in Fig. 5 demonstrate that despite the initial greedy pruning of candidate nodes the hybrid method obtains performances similar to direct optimal node selection based on OCS-GS. Since finding optimal node locations is a one-time exercise, the hybrid method offers a feasible alternative to find good node locations for constructing WESNs.

## VI. Conclusion

In this paper, we proposed an MIQP-based channel selection method which performs optimal channel selection for EEG in a least-squares based neural decoder design. We used this optimal channel selection method to investigate the optimality gap of state-of-the-art approximate channel selection methods compared to an optimal selection. We found the greedy method based on the LS-utility metric to perform similar to the optimal channel selection in an AAD task for standard cap EEG channels but requiring considerably less computation time, thereby providing a practical solution for the channel selection problem. We also used a topology-constrained modification of the MIQP to solve a WESN node selection problem with galvanic separation constraints. We showed that the galvanic separation constraints do not appear to have a significant impact on the AAD performance. Finally, to reduce the computation time but still include topological constraints and obtain near-optimal channel selection results, we proposed a hybrid approach of MIQP-based channel selection with greedy utility-based pruning, which showed no significant optimality gap with the optimization of the full MIQP.

## Footnotes

1 An evaluation involves training an optimal decoder for the given selection, and testing the performance of the resulting decoder.

2 An open-source MATLAB and Python based implementation of this method is available on https://github.com/mabhijithn/channelselect.

3 *M* should be chosen larger than the entry with the maximal absolute value in the final solution 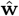, which is unknown before solving (7a)–(7d). In practice, if the magnitude of one of the entries in the final solution 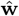 is equal or close to *M*, this means *M* has been set to a too small value, in which case the procedure has to be restarted with a larger *M*.

4 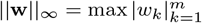where **w** = [*w*_1_ *…, w_m_*]^*T*^

5 In the experiments in this paper, we used the implementation of gLASSO from [26].

6 LME model based statistical analysis is not used in all of the comparisons in the remaining of this paper since the check for normality of the residuals failed in all comparisons, implying a bad fit of the model.

